# Pandemic Risk Assessment for a Swine Influenza A Virus in Comparative Human Substrates

**DOI:** 10.1101/2024.02.24.581872

**Authors:** Ian Padykula, Lambodhar Damodaran, Kelsey T. Young, Madelyn Krunkosky, Emily F. Griffin, James F. North, Peter J. Neasham, Vasilis C. Pliasas, Chris L. Siepker, James B. Stanton, Elizabeth W. Howerth, Justin Bahl, Constantinos S. Kyriakis, S. Mark Tompkins

## Abstract

Swine influenza A viruses pose a public health concern as novel and circulating strains occasionally spill over into human hosts, with the potential to cause disease. Crucial to preempting these events is the use of a threat assessment framework for human populations. However, established guidelines do not specify what animal models or *in vitro* substrates should be used. We completed an assessment of a contemporary swine influenza isolate, A/swine/GA/A27480/2019, using animal models and human cell substrates. Infection studies *in vivo* revealed high replicative ability and a pathogenic phenotype in the swine host with replication corresponding to a complementary study performed in swine primary respiratory epithelial cells. However, replication was limited in human primary cell substrates. This contrasted with our findings in the Calu-3 cell line, which demonstrated a replication profile on par with the 2009 pandemic H1N1 virus. These data suggest selection of models is important for meaningful risk assessment.

**Article Summary Line:** A novel swine influenza A virus was rigorously assessed for pandemic potential in animal models and human surrogate culture models, illustrating a wide range in potential public health risk dependent on the model utilized.

---

The ability of influenza A viruses of various genetic backgrounds to infect swine is of significant concern to public health. Swine have been proposed as key intermediate hosts in adaptation of avian influenza viruses to mammalian species, but they are also susceptible to infection with human influenza isolates. As multiple influenza viruses infect a single individual, they have the potential for reassortment of gene segments that can result in a novel virus to which the host has little or no immunity. This was exemplified in the 2009 H1N1 pandemic, in which gene segments from human, avian and swine origins reassorted to create a novel, antigenically shifted virus that rapidly spread across the globe (1, 2).

Currently endemic in swine populations are influenza strains belong to the H1N1, H1N2 and H3N2 subtypes (3). In the U.S., human seasonal influenza spilled into commercial swine populations in the early 2000s, reassorting with circulating swine influenza A viruses to create a lineage of H1N2 that possesses external proteins of human seasonal influenza origin with a swine internal gene cassette (4, 5). This lineage, designated 1B.2.2 or δ-2, has continued to diversify within swine populations since this point of introduction. This has resulted in an increasingly dominant endemic virus within commercial herds as well as several human cases of H1N2 variant (H1N2v) viruses (6). The ability of these swine viruses to spill back into human populations suggests there is limited immunity from cross-protective antibodies elicited by human seasonal influenza (7). As such, increased surveillance efforts have been focused on these viruses.

In order to properly assess the risk potential of emerging influenza strains, a framework has been established by the Center for Disease Control and Prevention, the Influenza Risk Assessment Tool (IRAT) (8). The tool allows for a systematic evaluation of emerging zoonotic influenza strains and a baseline comparison for prioritization and allocation of funding. By analyzing a given virus according to three main criteria, viral properties, host properties, and epidemiological factors, the tool seeks to answer two key questions. The first is that of emergence, that is, what is the risk of a novel influenza strain being capable of sustained human-to-human transmission. The second question is that of impact of viral infection on public health if sustained transmission is possible. The three main criteria of the IRAT are broken down into a total of 10 elements, several relating to a virus’s ability to transmit and cause disease within animal species as well as humans. The usage of animal models to assess risk to human populations is quite varied and involves additional cost and risk to investigators. Several *in vitro* model systems have also been used to determine susceptibility of human respiratory tissues, with increasingly complex culture systems allowing for growth and differentiation of primary epithelial cell cultures from host animals that approximate the *in vivo* environment (9-13).

We isolated an influenza A virus from a lethal case of swine influenza, A/swine/Georgia/A19-27480/2019 (H1N2; GA/19) that we then assessed in multiple models of infection to determine the extent of pathogenesis in its host species as well as the transmission potential to humans. Phylogenetic analysis determined the isolate as belonging to the 1B.2.2 lineage, with genetic relatedness to Midwest swine influenza viruses but also H1N2v viruses. Infections in mice, swine and ferrets showed an ability of the virus to infect multiple animal species and transmit to naïve contact animals. We further investigated risk to humans through infections in tissue culture systems, including Calu-3 cells and primary respiratory epithelial cells from human donors. Despite the human origins of both these substrates, results presented differing susceptibility of human tissues to the swine isolate that was dependent only on the model used.

## Materials and Methods

### Virus Isolation

A 6-month-old Hampshire cross market gilt was submitted for necropsy to the Athens Veterinary Diagnostic Laboratory, College of Veterinary Medicine, UGA after a sudden and severe illness of unknown etiology (14, 15). The animal was a 4-H show pig with a recent history of travel to an event. On necropsy, there was severe cranioventral consolidation with necrotizing bronchointerstitial pneumonia and ulcerative tracheitis; confirmatory fluorescent antibody testing and immunohistochemistry (IHC) on lung tissue was diagnostic for influenza A bronchitis. Bronchiolar sections of lung tissue weighing approximately 100 grams were homogenized and the resulting homogenate passed through a 0.45 µm nylon mesh filter. The homogenate was then used to inoculate flasks of MDCK cells. The virus was passaged twice blindly, then plaque purified. The plaque-purified virus was used in further animal and *in vitro* infection studies.

### Phylogenetic Analysis

Viral sequence data was generated using a MinION platform as previously described (15). MinION reads were assembled using IRMA v 0.6.7 (16) and validated by Illumina sequencing as previously described (17). Sequences are publicly available in GenBank, under BioProject PRJNA600894, NCBI accession number (pending). To investigate the potential for the isolates involvement in human-swine transmission and the virus’s evolutionary history the consensus sequences for the HA protein (H1) and NA protein (N2) were used in phylogenetic analyses. Nucleotide sequence data was collected from the NCBI Influenza Virus Resource (date range 6/6/2014 through 7/2/2019) and grouped into separate datasets for human H1, swine H1, human N2, swine N2, human M, and swine M (18).

To determine potential human-swine transmission associated with the swine isolate A/swine/Georgia/A19-27480/2019 (H1N2), maximum likelihood trees were created by performing 100 bootstrap replicates of RAxML v 8.2.4 for the coding regions of each segment dataset (19). The trees were created using a generalized time reversible nucleotide substitution model with gamma distributed rate variation among sites (GTR+ Γ4). A root-to-tip regression of the estimated trees was performed to determine the molecular clock signal of the data and temporal outliers were identified and removed using TempEst v 1.5.3 (20).

Bayesian phylogenetic tree estimation was performed for a randomly sampled subset of n = 450 taxa for the swine HA, and NA segment datasets using BEAST v1.10.4 (21). Six independent Markov Chain Monte Carlo (MCMC) runs for both proteins were preformed using a GTR+ Γ4 substitution model, lognormal uncorrelated relaxed clock model, and a Gaussian Markov Random Field Skyride coalescent (22, 23). Each MCMC run had a chain length of 100 million states, sampling every 10 thousand states. After removing appropriate burn-in from the beginning of the run (10%) a maximum clade credibility phylogenetic tree was generated for each segment from a posterior sampling of 9000 trees.

### Animal Infection and Transmission Experiments

Five- to eight-week-old female DBA/2 and BALB/c mice (Jackson Labs and Envigo, respectively; n=25 per strain) were divided into groups, infected (n=20) and mock (n=5). Mice were infected under isoflurane anesthesia via intranasal inoculation with 1e5 pfu of GA/19 virus in a 30 µL volume. Mock infected mice were given 30 µL PBS intranasally. Animals were observed for clinical signs twice daily and weighed daily. At 2- and 4-days post-infection (dpi), a subset of 5 mice from each infected group was euthanized and lungs collected for virus titration. At 5 dpi, 3 mice from each infected group were euthanized and lungs collected and perfused with neutral-buffered 10% formalin before submission for histopathological examination. The remaining mice were weighed daily until 13 dpi. All mice were euthanized at 25 dpi and serum samples collected.

Six, 6-week-old, influenza virus-naïve and porcine reproductive and respiratory syndrome virus (PRRSV)-naive conventional cross-bred Yorkshire/Hampshire male and female pigs were obtained from Auburn University’s Swine Research Center (an influenza virus and PRRSV-seronegative herd), were obtained for the swine study. One week prior to infection the study animals were treated with ceftiofur crystalline free acid (Zoetis). One day before infection animals were sedated with an intramuscular injection of ketamine (0.5 mg/kg), xylazine (0.5 mg/kg) and tiletamine-zolazepam (1 mg/kg) and baseline bronchoalveolar lavage (BAL) samples and blood samples taken. Animals were then separated into two groups, infected (n=3) and contact (n=3). Infected animals were inoculated intranasally with 1 ml of 1e6 pfu/mL GA/19 virus in each nostril. Temperatures and nasal swabs were taken daily after infection. At 2, 4 and 6 dpi infected animals were sedated and BAL and blood samples were collected. At 3 dpi contact animals were co-housed with infected animals. Nasal swabs were performed daily on infected animals and on contact animals after introduction. At 13 dpi blood samples were collected from all animals.

Ten 12-week-old ferrets were divided into two groups, infected (n=6) and contact animals (n=4), with equal gender distribution between the two groups. Three days prior to infection, all animals were anesthetized under isoflurane and 3 mL venous blood draws performed, as well as placement of subdermal temperature transponders (BMDS) for animal identification and temperature monitoring. On day 0, six animals were anesthetized and inoculated intranasally with 1e6 pfu of GA/19 virus in a 1 mL volume distributed equally between each naris. Nasal washes and weights were taken on infected animals one day post-infection and every other day subsequent. At 2 dpi, naïve contact animals were co-housed with infected animals in a 1:1 ratio and nasal washes and weighing performed as on infected animals. At 4 dpi, two infected animals were euthanized and tissue samples collected from the respiratory tract. At 7 and 14 dpi venous blood draws were performed, with animals euthanized at 14 dpi.

All animal studies were reviewed and approved by the Institutional Animal Care and Use Committee (IACUC) prior to initiation.

### Cell Culture and Infection

Calu-3 (ATCC), normal human bronchial epithelial (Lonza), and porcine primary nasal epithelial (24) cells were cultured and differentiated at an air-liquid interface (ALI) in 12-well plates as previously described (9, 10). In brief, cells were seeded onto collagen-coated transwells of a 12-well plate (Corning). Media was changed 1 day after seeding and every other day afterward until cells had reached confluency. Apical media was then removed from the transwell and basolateral media replaced with ALI media (DMEM/F-12 supplemented with 1% penicillin/streptomycin, 2% NuSerum and 50 nM retinoic acid). At day 0 apical surfaces were washed and inoculated with GA/19 virus at an MOI of 0.01 in a 200 µL volume. All infections were run in triplicate wells. Cultures were then incubated in a humidified 5% CO_2_ incubator at 37° C for 2 hours before inoculum was removed. At 12, 24, 48, 72, and 96 hours post-inoculation, 1 mL of sterile PBS was used to wash the apical cell surface, then the wash was assayed for viral titers.

### Sample Processing

Mouse lungs were individually placed in 1 mL PBS after collection and kept on ice until homogenization. Lungs were homogenized as previously described (25) clarified by centrifugation, aliquoted, and then stored at −80° C. Bronchioalveolar lavage (BAL) fluid was placed on ice immediately after collection and then passed through a 40 micron filter before aliquoting and freezing at −80° C. Nasal swabs were collected using polyester swabs, placed in tubes containing 2.0 ml of phosphate-buffered saline (PBS) supplemented with 1% antibiotic-antimycotic, and placed on ice immediately after collection. Samples were then sonicated for 10 minutes before aliquoting and storage at −80° C (26). Nasal washes were performed with sterile PBS, then samples were placed on ice after collection, clarified, aliquoted, and then stored at - 80° C.

### Virus Titration

For plaque assays, tenfold serial dilutions were prepared from thawed samples in serum-free MEM. An inoculum of 250 µL was placed into each well of a 12-well plate of confluent Madin-Darby canine kidney (MDCK) cells (MDCK-ATL, FR-926, International Reagent Resource) and incubated in a humidified chamber at 37° C for 1 hour before a 1.2% Avicel overlay supplemented with 1x TPCK trypsin was placed on the culture. Plates were then incubated for 72 hours in a 5% CO_2_ humidified chamber at 37° C before the overlay was removed and plates fixed with an 80% methanol 20% acetone solution. Fixed wells were then stained with crystal violet to visualize plaques that were counted, and then infectious virus titers calculated.

For virus quantitation by PCR, viral RNA was extracted from nasal swab samples and tested for the matrix gene of IAV. Viral RNA was extracted using RNAzol RT. RT-PCR was performed using Taqman® Fast Virus 1-Step Master Mix. A 25 µL PCR mixture containing 6.25 µL 4X Fast Virus Master Mix, 14.75 µL DNase/RNase-free distilled water, 0.5 µL of each primer (forward: AGATGAGTC TTCTAACCGAGGTCG, reverse: TGCAAAAACATCTTCAAGTCTCTG), 1 µL of probe (FAM-TCAGGCCCCCTC AAAGCCGA-BHQ), and 2 µL of sample RNA template was prepared. Reactions were run at 50° C for 30 minutes, followed by 95° C for 15 minutes, followed by 40 cycles at 95° C for 10 seconds then 60° C for 20 seconds. Data were acquired on the BioRad C1000 Touch Thermal Cycler and data analysis performed with BioRad CFX Manager (v3.1). Viral titers were calculated based upon titration of a stock of known concentration, given as relative expression units (REU) compared to a negative control sample.

### Statistical analysis

Viral titers were analyzed using analysis of variance (ANOVA) using GraphPad Prism version 8.0.0 for Windows (GraphPad Software, San Diego, California USA, www.graphpad.com). A p-value ≤ 0.05 was considered significant.

## Results

### Phylogenetic assessment

A purified viral stock was obtained from lung tissue of a diseased pig and then plaque purified before sequencing using a MinION platform. Obtained consensus sequence data identified the virus as belonging to the H1N2 subtype (1B.2.1) (15) and was confirmed by subsequent Illumina deep sequencing. We performed phylogenetic analysis of the isolate in the context of human and swine H1N2 viruses to determine genetic distance from recent human influenza viruses of similar genetic backgrounds. Maximum-likelihood trees assembled from sequences extending five years prior to the date of sample collection revealed a significant distance between the isolate and the closest human isolate (Supplementary Figure 1). However, the isolate did demonstrate a close relationship to H1N2 variant (H1N2v) cases in which a swine influenza virus resulted in limited human infections (Figure 1, Figure 2) (27, 28).

**Figure 1.**
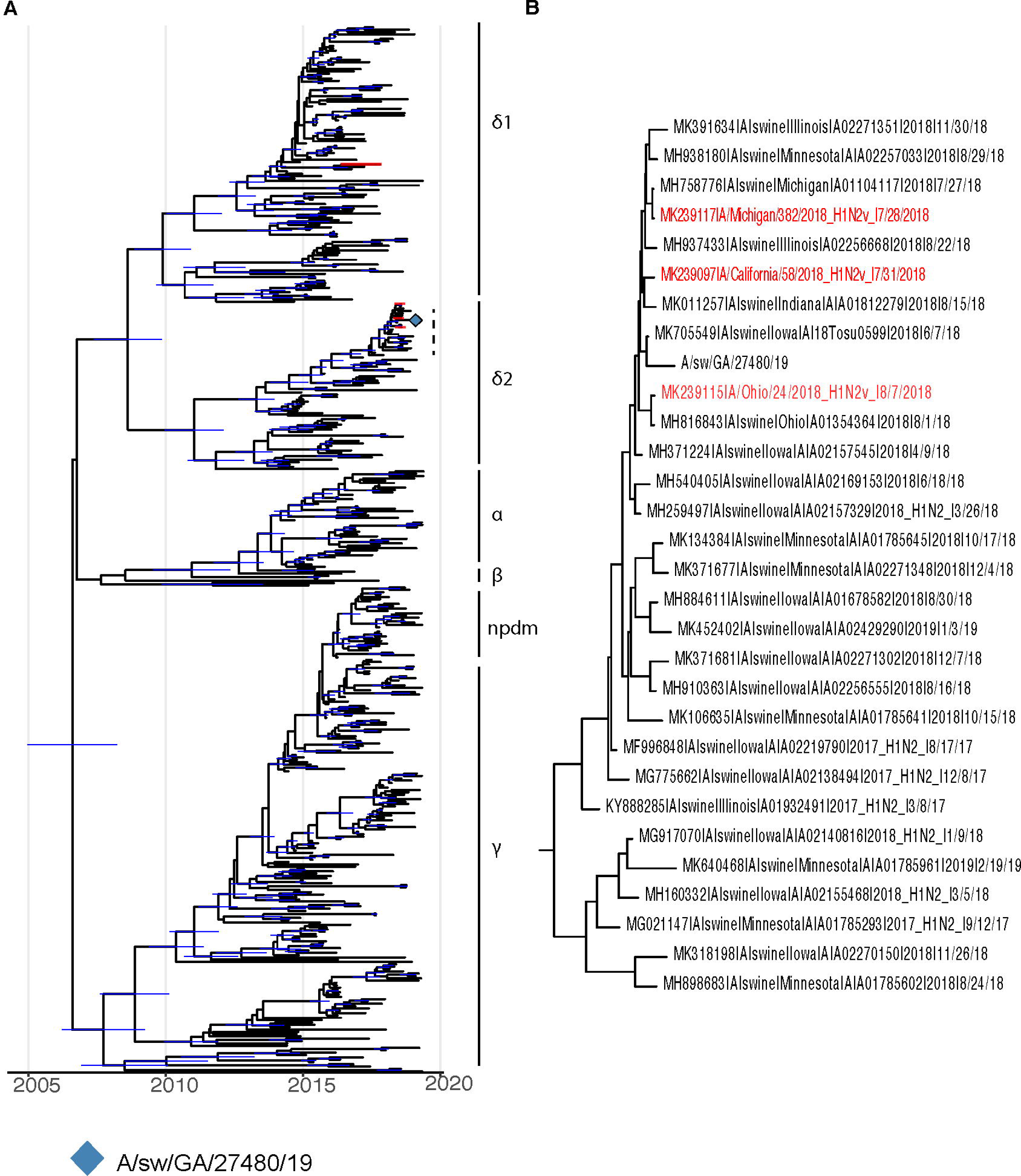
BEAST phylogenies for swine influenza A H1Nx isolates collected from 2014 to 2019. A, B) Phylogenetic reconstruction for hemagglutinin (HA) segment of swine H1Nx isolates. Nodes with a posterior support of greater than 95% are annotated with a 95% Bayesian Credible Interval in blue. Taxa for H1N2 variant isolates are colored in red.

**Figure 2.**
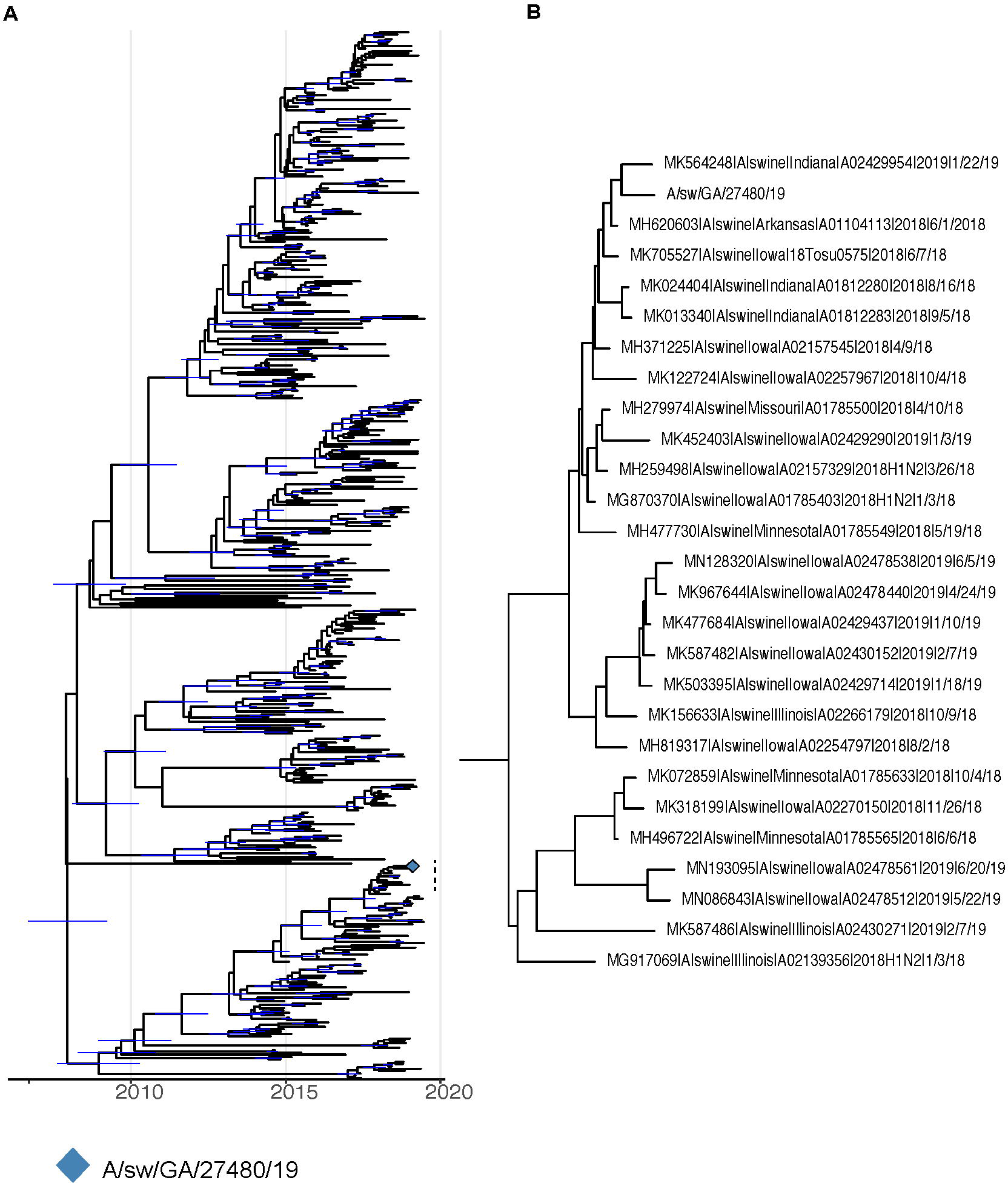
BEAST phylogenies for swine influenza A HxN2 isolates collected from 2014 to 2019. A, B) Phylogenetic reconstruction for neuraminidase (NA) segment of swine HxN2 isolates. Nodes with a posterior support of greater than 95% are annotated with a 95% Bayesian Credible Interval in blue. Taxa for H1N2 variant isolates are colored in red.

Bayesian phylogenetic trees were created for the HA (Figure 1a-b) and NA (Figure 2a-b) genetic segments. The resulting trees illustrate a close genetic distance between the GA/19 isolate and contemporary swine influenza viruses circulating in the Midwest United States. Interestingly, while phylogenies among the gene segments reveal a genetically drifted variant from otherwise unremarkable swine influenza isolates, they do not explain the virulence of the initial case presentation from which the isolate was obtained.

### Viral replication in the murine model

Although mice do not shed influenza virus or exhibit symptomology correlating to swine or human disease, virulence of influenza infection in mice has been shown to correlate to severity of disease in the case of several human and swine influenza strains (15, 25, 27). We assessed virus replication, pathogenesis, and clinical disease in BALB/c and DBA/2 mouse strains as both are commonly used to measure influenza disease and DBA/2 are notably susceptible to influenza virus infection, often displaying increased disease as compared to other mouse strains (17, 29, 30). Intranasal infection with the GA/19 strain of swine influenza did not cause weight loss in either DBA/2 or BALB/c mice, although there were differences in weights between infected and naïve mice from each background, particularly among mice belonging to the DBA/2 group (Figure 3a). Despite the lack of weight loss, there was robust viral replication at 2 days post infection, with average lung titers from DBA/2 mice of 5.28e5 pfu/mL and BALB/c mice at 2.64e4 pfu/mL of lung homogenate. On day 4, titers had reduced significantly for DBA/2 mice to 8.32e4 pfu/mL, while BALB/c mice virus titers, at 2.74e4 pfu/mL, were not reduced (Figure 3b). At both timepoints DBA/2 mice had significantly higher lung titers than BALB/c animals assayed at the same time. Pathological findings on day 5 showed mild pathology in lung tissues, regardless of genetic background. Interestingly, DBA/2 mice had a noted increased degree of necrotic bronchiolar epithelium and lymphocytic migration around vessels compared to BALB/c mice (data not shown). These results are consistent with previous studies demonstrating the enhanced susceptibility of DBA/2 mice to influenza infection (17, 29, 30).

**Figure 3.**
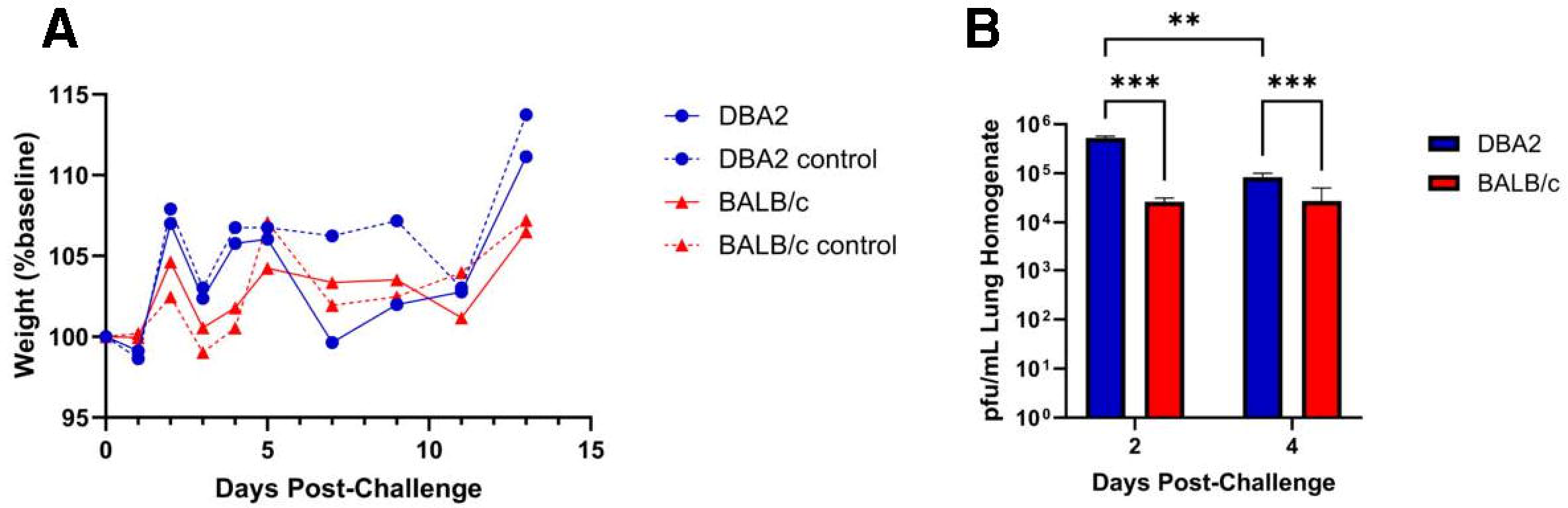
Weight change and lung virus replication following infection of DBA/2 or BALB/c mice with A/sw/GA/27480/19 (H1N2). Five- to eight-week-old BALB/c and DBA/2 mice (n=25 mice/group) were inoculated intranasally with either 1e5 pfu of virus in a 30 µL volume (n=20) or with PBS (n=5). Weight loss was tracked for 13 days post infection (dpi) (a), dashed lines represent mock-infected control groups. At 2 and 4 dpi, a subset of 5 mice from each infected group were euthanized and lungs collected. Viral titers, described a pfu/mL of lung homogenate, were determined by plaque assay (b). Significance values of ≤0.005 and ≤0.0005 are denoted by ** and ***, respectively. Error bars indicate mean ± SD.

### Viral replication and transmission in swine

The GA/19 virus was isolated from a Hampshire gilt, 4-H show pig that developed a fever and died suddenly after travel to an event (14, 15), so we sought to assess GA/19 infection in healthy pigs. Clinical symptoms in infected pigs were mild, as seen in experimental infections with other swine influenza viruses following intranasal inoculation (31, 32). Viral titers in BAL fluid averaged 2.00e5 pfu/mL at day 2 post-infection, decreasing slightly at day 4 post-infection to 1.56e5 pfu/mL (Figure 4). The virus transmitted to 2 out of 3 contact pigs, as seen by shedding in nasal swabs (Figure 4). Three days after co-housing, nasal swabs from contact animals averaged 7.94e1 pfu/mL and remained positive for viral shedding until day 6 post-contact. Nasal swab samples for infected and contact animals were also assayed for influenza virus load by qPCR, which showed REU levels corresponding to BAL titers among infected individuals (Supplementary Figure 2). However, by this assay all contact animals became positive by 3 days post-contact. All animals were positive for infection as determined by serology, as well (data not shown). Despite rapid recovery, contact animals showed mild symptoms, limited to lethargy and elevated temperature (data not shown).

**Figure 4.**
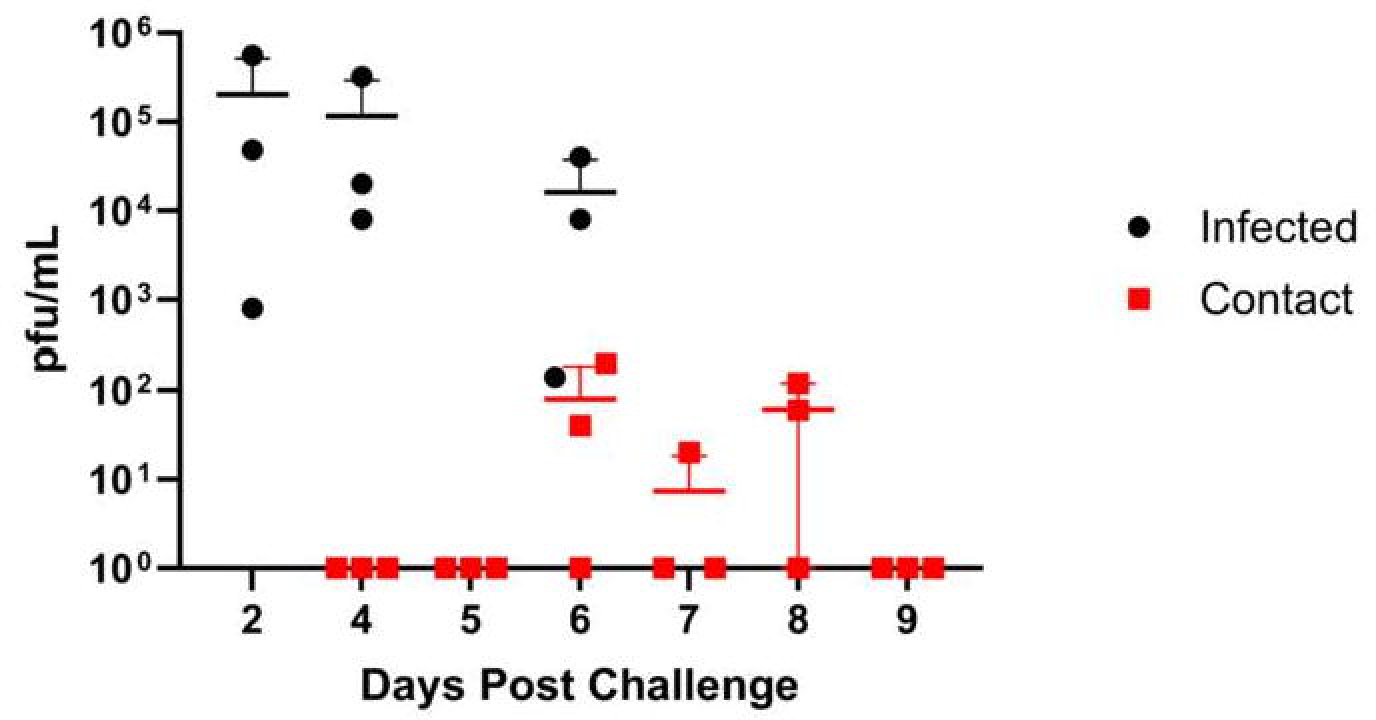
Replication and transmission of A/sw/GA/27480/19 (H1N2) in a swine model. Six-week-old pigs were inoculated intranasally with 2e6 pfu of GA/19 in a 2 mL volume (n=3). At 2, 4 and 6 dpi BAL samples were collected (black data points). At 3 dpi, naïve contact animals (n=3) were co-housed with infected animals and nasal swabs collected daily (red data points). Viral titers were determined by plaque assay. Error bars indicate mean ± SD.

### Viral replication and transmission in ferrets

As ferrets are considered the “gold standard” animal model for human influenza virus infection (33), we assessed infection and clinical disease in this model. Ferrets infected with GA/19 showed greatest viral titers in nasal washes at 1 dpi, with most animals clearing virus by day 5, with one animal remaining positive at this timepoint (Figure 5). Infection resulted in weight loss peaking at 3 dpi, however this was mild with average weights above baseline by 9 dpi (Supplementary Figure 3). Contact ferrets introduced at 2 dpi were all positive for viral shedding by 5 dpi (day 3 post-contact (dpc)). Contact animals displayed a pattern of viral shedding similar to infected animals (Figure 5), with all animals negative in nasal washes by 11 dpi (8 dpc). Weight loss within the contact group was also mild, but remained depressed through 11 dpi.

**Figure 5.**
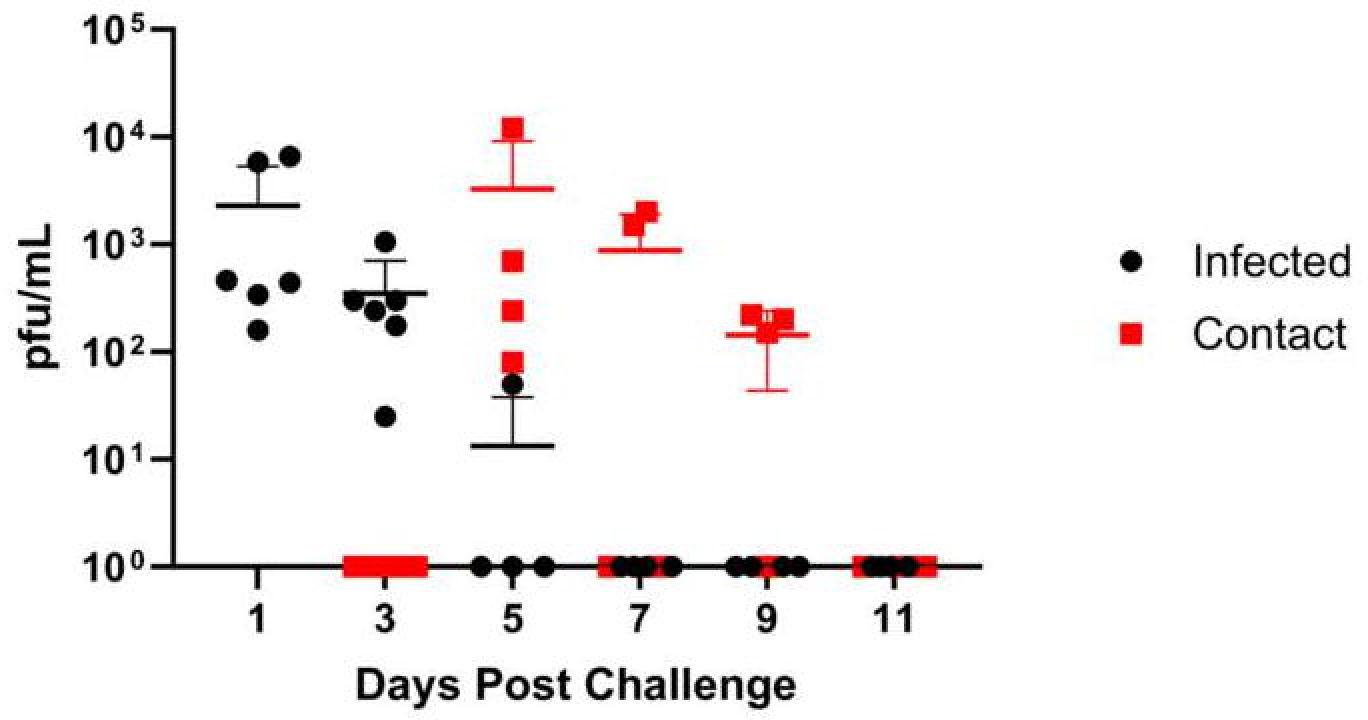
Replication and transmission of A/sw/GA/27480/19 (H1N2) in ferrets. Twelve-week-old ferrets were inoculated intranasally with 1e6 pfu of GA/19 in a 1 mL volume (n=6). At 1, 3, 5, 7, 9 and 11 dpi, nasal wash samples were collected and titers evaluated by plaque assay (black data points). At 2 dpi, naïve contact animals (n=4) were co-housed with infected animals (1:1) and nasal washes taken and evaluated for virus titer by plaque assay (red data points). Error bars indicate mean ± SD.

### Viral replication in *in vitro* substrates

To further investigate capability of GA/19 to replicate in human respiratory tissues we cultured Calu-3 cells, normal human bronchial epithelial (NHBE) cells, and as a control, porcine nasal epithelial (PNE) cells at an air-liquid interface (ALI). Upon differentiation and culture at ALI, all three cell substrates produced mucus and reflected the airway surface. The airway cell substrates were infected on the apical surface with GA/19 (MOI of 0.01) and the apical surface sampled at 12 and then every 24 hours to determine virus replication and differing levels of permissiveness to infection. Infection of PNE cells (24) resulted in the highest titers of GA/19, rapidly increasing to 1.62e6 pfu/ml at 24 hours post-inoculation and then peaking at 3.98e7 pfu/mL by 48 hours post-inoculation (Figure 6a). Calu-3 cells, derived from a human lung adenocarcinoma, are frequently used as a model of susceptibility to viral infection (34, 35). Our infections with the Calu-3 substrate with GA/19 showed rapid viral replication, reaching peak titers of 2.34e5 pfu/mL by 48 hours post-inoculation (Figure 6a). Surprisingly, infection of NHBE cells with the swine isolate showed contrasting results to the other cell substrates. Viral titer remained depressed compared to Calu-3 and PNE substrates at equivalent timepoints, never exceeding 1e3 pfu/mL. Timing of peak viral titer was also retarded, being seen at 72 hours post-inoculation as opposed to 48 hours observed for PNE and Calu-3 cells (Figure 6a). GA/19 virus replication kinetics were confirmed in Calu-3 and NHBE cells of a different donor background with a duplicate experiment that demonstrated near identical virus titers over the 96-hour assay (Figure 6a, dashed lines). While there was little to no virus replication, GA/19 was able to infect as demonstrated by confocal imaging of virus NP antigen staining of NHBE cells at 96 hours post-inoculation (Supplementary Figure 4).

**Figure 6.**
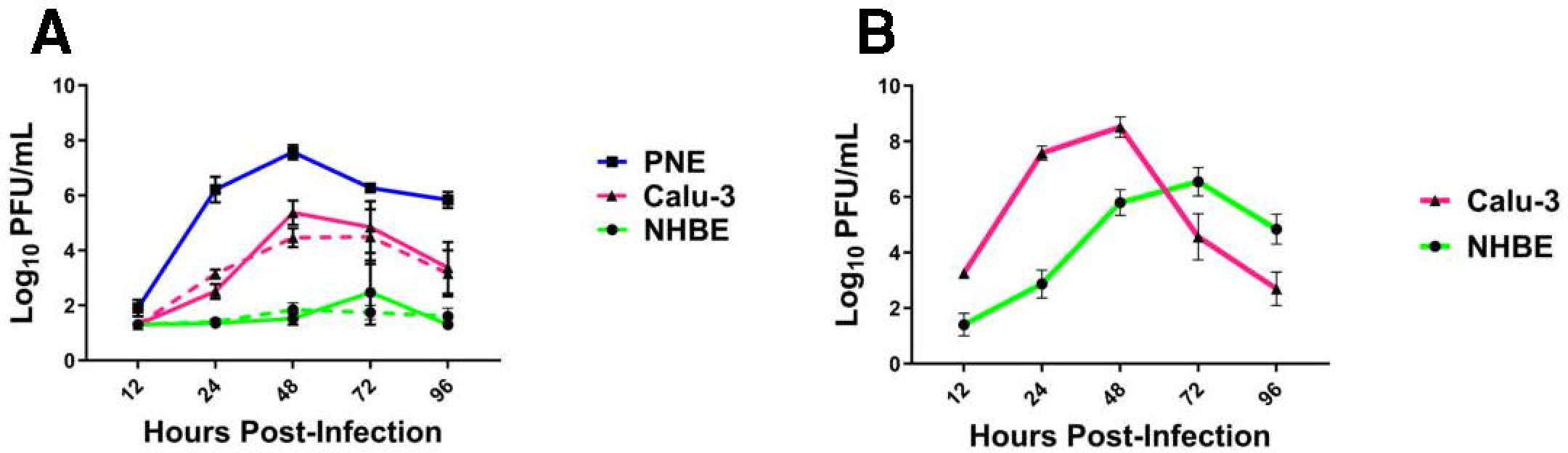
Replication of H1 influenza viruses *in vitro*. Primary porcine nasal epithelial (PNE), primary human bronchial epithelial (NHBE), and Calu-3 cells (all at ALI) were infected apically with either GA/19 (A) or A/CA/07/09 (B) at an MOI of 0.01. At 12, 24, 48, 72, and 96 hours post-infection the apical surface of cultures were washed, the fluid collected, and then titered for virus by plaque assay. Dashed lines denote duplicate experiments. Error bars indicate mean ± SD.

To confirm the permissiveness of the human cell substrates to a human influenza virus we ran a complementary infection experiment with the 2009 pandemic virus, A/CA/07/09 (CA/09; MOI of 0.01). Calu-3 cells infected with CA/09 showed a pattern of replication kinetics markedly similar to that seen with GA/19, peaking at 48 hours post-infection at 3.24e8 pfu/mL (Figure 6b). NHBE cells infected with CA/09 also displayed a similar kinetics pattern to infections with GA/19, peaking at 72 hours, but resulted in much greater titers at 24, 48, 72, and 96 hours post-inoculation and a peak virus titer of 3.47e6 pfu/mL by 72 hours post-infection (Figure 6b) compared to the minimal levels seen with the swine-origin virus.

## Discussion

Our studies show a notable discrepancy in outcomes of infection with the spectrum of models used to assess pandemic risk of zoonotic influenza viruses. Utilizing specific elements within each of the three main criteria of the IRAT tool, we assessed the potential of the A/swine/Georgia/A19-27480/2019 (GA/19) virus to impact human health. To investigate viral characteristics we performed a phylogenetic analysis of sequences of both the HA and NA genes. This analysis demonstrated a close relationship of the GA/19 virus to contemporary swine H1-δ2 viruses within North America, as well as a surprisingly close relationship to human variant influenza isolates. Our studies further examined host interactions with the virus utilizing several animal models to assess pathogenicity as well as transmission potential.

Every animal model of infection used showed unequivocal replication of the GA/19 virus, as well as transmission potential in both swine and ferret hosts. Corroborating these results are infection experiments in human and swine-derived airway epithelial cell culture models. The virus displayed similar growth kinetics in human Calu-3 and porcine nasal epithelial cells, peaking in both systems by 72 hours post-infection albeit with differing peak titers. Contradictory to these findings were the results of replication kinetics experiments in NHBE cells, a widely used model to ascertain permissivity of human cells to viral infection (36). Within this latter culture model, infection with the GA/19 virus resulted in little to no viral replication. Despite the lack of replication, confocal images confirmed viral invasion into the cell substrate by 96 hours post-infection suggesting the NHBEs were susceptible to infection, but the virus failed to replicate. Infection of the same NHBE cells with a human-origin A/CA/07/2009 (pdmH1N1) confirmed them as permissive to IAV infection. Contrary to initial infections in NHBE cells, a repeat experiment using an alternate donor with varying demographics showed a modest capacity of the GA/19 virus for replication in human primary respiratory epithelial cells (Supplementary Figure 5). The inter-donor variability of NHBE cells is a recognized problem in the context of influenza infection; however, the cause of this is likely multifactorial. Recent studies have demonstrated differential kinetics and amplitude of interferon gene expression subsequent to infection of NHBEs with human-origin H1N1 and H3N2 viruses. However, proteomic studies indicate that the problem may be even more nuanced. Here the expression of proteins critical to control of influenza virus replication, namely IFIT1, IFIT2, IFITM1, IFITM2, IFITM3 and MX, could have expression levels that varied as much as 10-fold between donors (37).

Of those animal models used in our studies, ferrets in particular are considered a gold standard for influenza A virus infection, virulence, and transmissibility in humans, and are thus used in assays to determine the risks posed by avian, porcine and emerging influenza A viruses (33, 38-40). The cellular ligand used by the influenza virus hemagglutinin to bind to and to enter human cells, an α2,6-linked sialic acid, shows similar distributions in ferret and human respiratory tissues (38, 41). Additionally, influenza infection in ferrets has a clinical disease phenotype very similar to what is seen in humans, including weight loss, sneezing, and lethargy (40). Our infection studies in the ferret model provided clear evidence of replication and transmission, a strong indication that the isolate would pose a threat to a human host.

Two different human-origin cell culture systems were used in our *in vitro* infections. Calu-3 cells are a continuous human epithelial lung cell line derived from a pulmonary adenocarcinoma, and have been characterized extensively and determined to display a sialic acid receptor profile that allows for infection with human influenza virus isolates (34, 35). Normal human bronchial epithelial cells are primary cells isolated from tracheobronchial tissue sections and have been used extensively in influenza research (36, 41). When cultured at an air-liquid interface these cells display a mixed morphology, including ciliated and mucus-secreting goblet cells. Importantly, these cells share the same α2,6 sialic acid receptor profile as Calu-3 cells, a critical component in mammalian influenza virus entry to the host cell. Despite these similarities, our infection studies showed strikingly different results between the two substrates. This hints at a subtler set of factors distinguishing the two cell substrates, creating a more permissive environment in Calu-3 cells for replication of a swine-origin influenza virus. While both models serve as useful tools in studying influenza infection in human respiratory tissues, this disparity must be taken into consideration for future risk assessments of emerging swine influenza isolates.

There were several limitations present in our study. First, in our ferret infection studies transmission was determined by placing naïve animals in direct contact with infected animals. More sophisticated housing systems have been created that would allow for determination of potential virus transmission by airborne droplets in addition to contact transmission. Also, NHBE susceptibility to infection varied, likely based upon donor variability. The extent of this variability is not known and would require extensive screening to estimate the consistency of these substrates. Finally, our *in vitro* infections used primary human cells isolated from a single area of the respiratory tract. While the influenza virus is normally capable of infecting cells throughout the respiratory tract, the process normally begins in the nasal cavity and upper trachea. Including these tissues in future studies would give a greater insight into tissue-specific host restriction of swine influenza viruses.

The GA/19 virus was isolated from a 6-month-old show pig that developed a high fever and died suddenly after recent travel to an event (14, 15). However, in our infection study with high-health status 6-week-old pigs, we observed only mild clinical signs despite robust replication in the lower and upper respiratory tract and efficient transmission to naïve contacts. Histopathological findings from the dead gilt found influenza A antigen in the lung and lymphoid tissue, but bacterial infection (*Streptococcus suis*) was also detected in the lung. Also, other viral infections (e.g. porcine circovirus, porcine reproductive and respiratory syndrome virus) were not excluded as predisposing factors (14). Nonetheless, influenza A virus infection was determined to be the primary diagnosis and the GA/19 isolate was highly related to other H1N2 virus infections across the Midwest, and two human H1N2v infections.

In summary, we have confirmed the ability of GA/19 to replicate and transmit in multiple models of human influenza virus infection. Among these, our infection studies in mice and ferrets agree with the literature in their utility as models of infection with swine influenza A viruses. We have also characterized the extent to which the isolate replicates in swine, its native host species. Importantly, our experiments we have highlighted considerations that must be taken in assessing pandemic risk of influenza strains *in vitro*, as commonly used models of human respiratory infections possess widely differing abilities to host zoonotic influenza viruses.

## Supporting information

Padykula et al. (2024) Supplementary Figures

## Acknowledgments

This project was funded in part with Federal funds from the National Institute of Allergy and Infectious Diseases, National Institutes of Health, Department of Health and Human Services under Contract No. HHSN272201400004C (NIAID Centers of Excellence for Influenza Research and Surveillance, CEIRS). The funding bodies had no role in the conceptualization, synthesis, or decision to submit this work for publication. Madin-Darby Canine Kidney (MDCK-ATL) Cells, FR-926, were obtained through the International Reagent Resource, Influenza Division, World Health Organization Collaborating Center for Surveillance, Epidemiology and Control of Influenza, Centers for Disease Control and Prevention, Atlanta, Georgia, USA.

## About the Author

Dr. Padykula is a postdoctoral scholar at the Medical University of South Carolina, Charleston, South Carolina. His research interests include identification, characterization, and vaccine development for emerging respiratory pathogens.

## Appendices

Padykula *et al.* (2024) Supplementary Figures

