## Supplementary material for "Pandemic Risk Assessment for a Swine Influenza A Virus in Comparative Human Substrates": Padykula et al. (2024) Supplementary Figures

**Supplementary Figure 1**

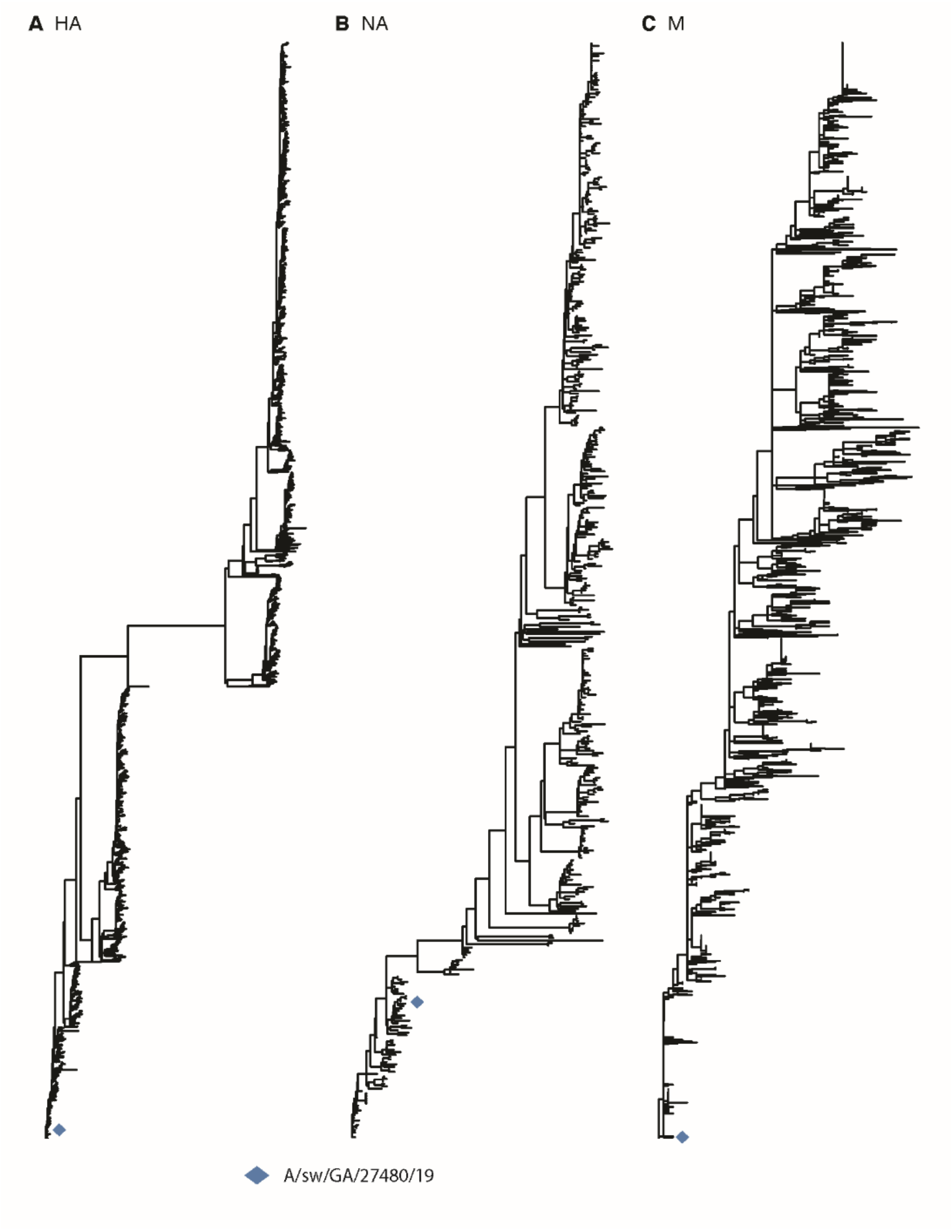


Supplementary Figure 1. Maximum likelihood phylogeny for swine isolates collected between 2014 and 2019. Phylogenetic reconstructions for the hemagglutinin (A), neuraminidase (B) and matrix (C) genes within the context of U.S. swine influenza A isolates representing the 5 year period prior to isolation of the GA/19 virus.

Supplementary Figure 2

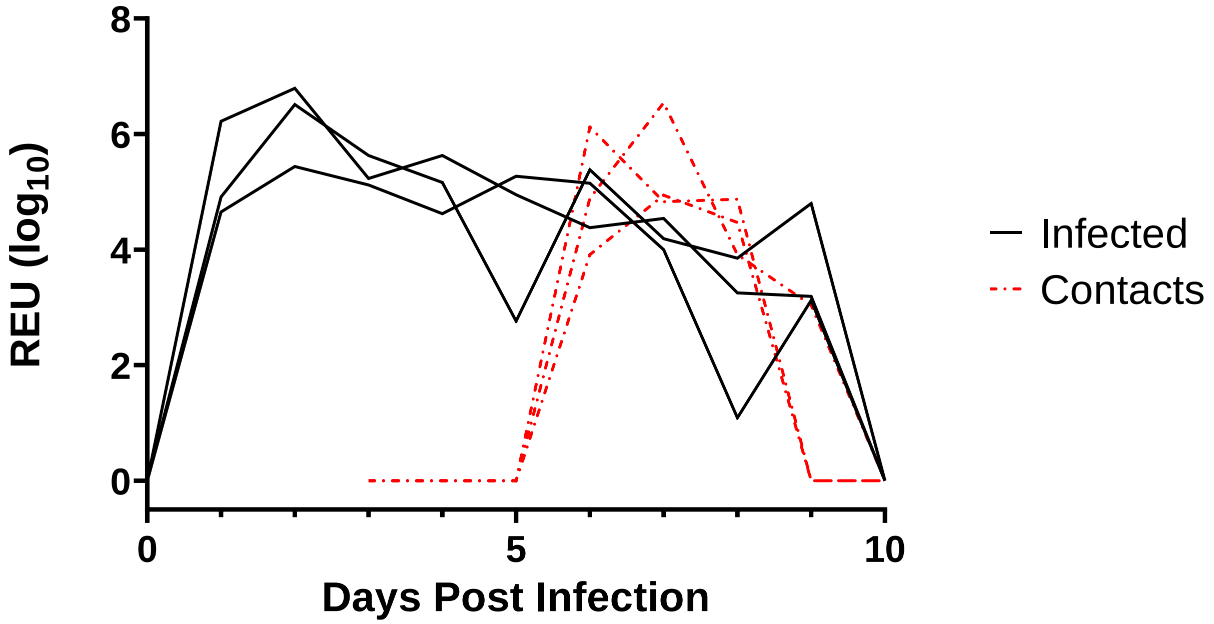


Supplementary Figure 2. Nasal shedding of A/sw/GA/27480/19 (H1N2) in swine as determined by qPCR. Infected animals showed peak viral loads by 2 dpi, remaining positive until day 9. All three contact animals became positive by 3 dpi, two remaining positive until day 8 and one until day 9.

Supplementary Figure 3

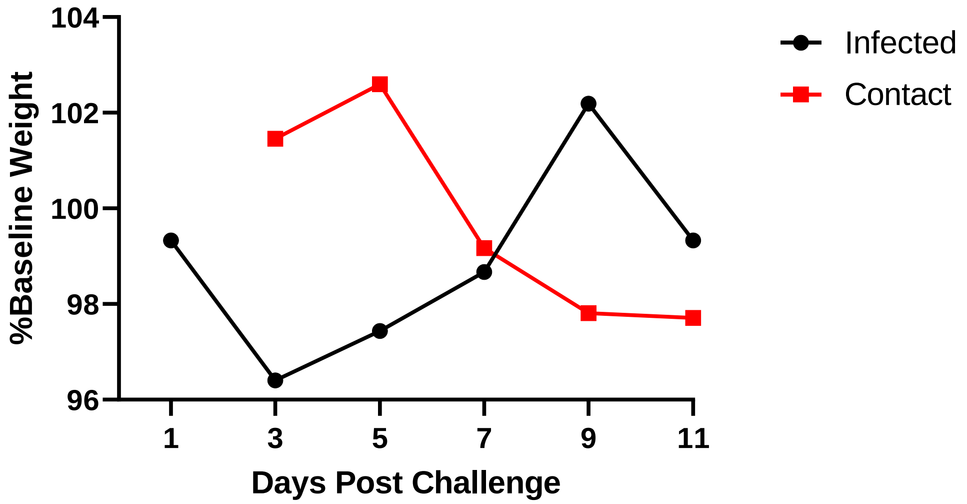


Supplementary Figure 3. Weight loss in ferrets post-challenge with A/sw/GA/27480/19 (H1N2). Weight measurements were taken until 11 dpi. Infected animals experienced an initial decrease in weight before returning to baseline levels. Contact animals experienced a comparable decrease in weight corresponding to an onset of viral shedding.

Supplementary Figure 4

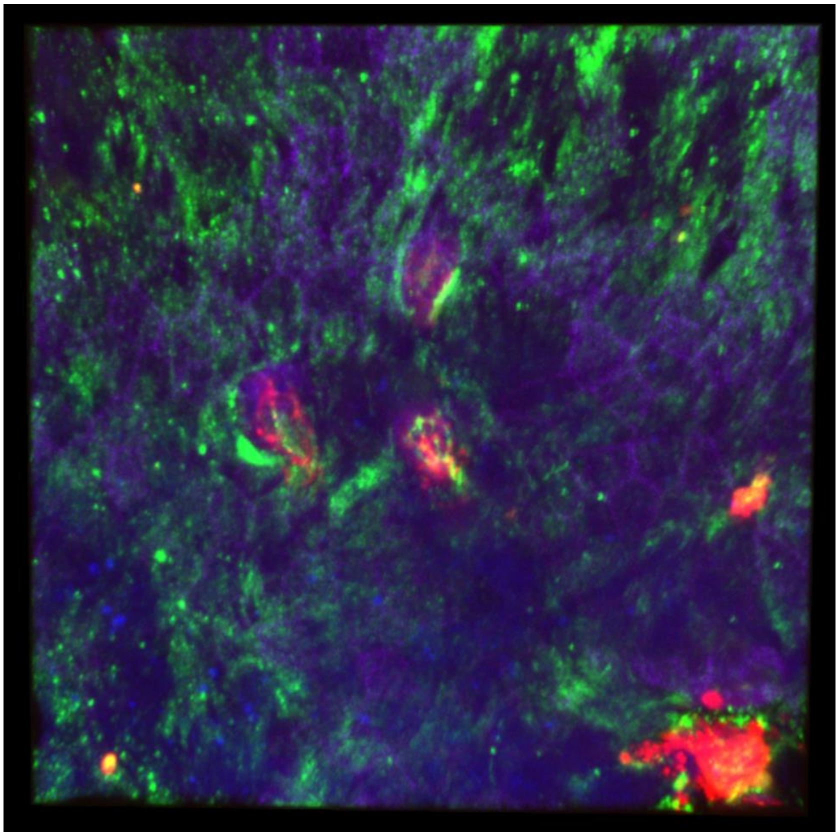

Supplementary Figure 4. Confocal image of NHBE cells infected with GA/19 at 96 hours post-infection. Despite showing minimal viral replication, NHBE cultures infected with GA/19 showed clear evidence of viral invasion. Red foci indicate cells infected with the swine influenza isolate. Red: viral nucleoprotein, blue: nuclear stain, purple: F-actin, green: beta tubulin.

Supplementary Figure 5

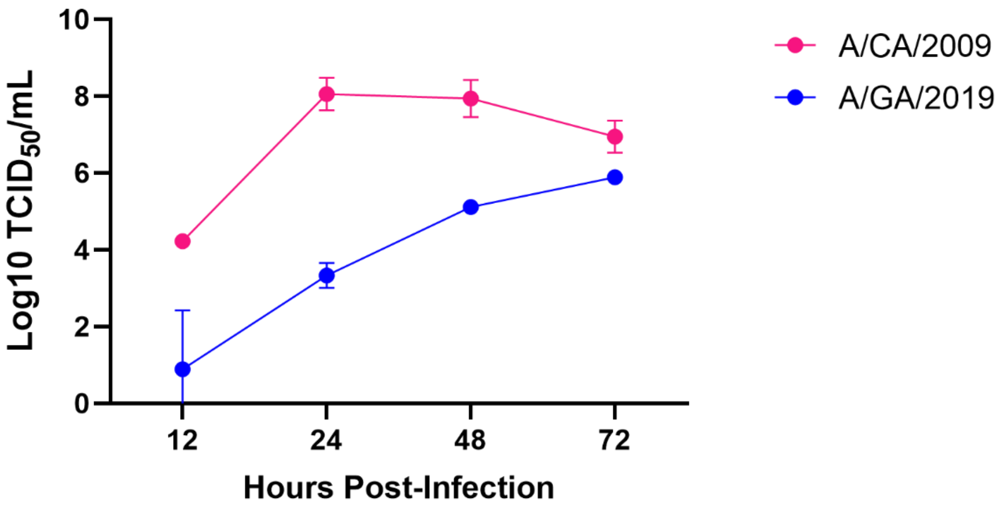


| Experiment | Donor Sex | Donor Age | Donor Race | Doubling Time (hours) |
|---|---|---|---|---|
| 1 | M | 56 | C | 27 |
| 2 | F | 66 | B & H | 31 |

Supplementary Figure 5. Replication kinetics of GA/19 compared to CA/09 in an NHBE cells from an alternate donor. Cultures were infected apically with either GA/19 or CA/09 at an MOI of 0.001. At 12, 24, 48, and 72 hours post-infection the apical surface of cultures were washed and the fluid titered for virus by TCID50. Error bars indicate mean ± SD.
